# Architectural Principles for Characterizing the Performance of Sequestration Feedback Networks

**DOI:** 10.1101/428300

**Authors:** Noah Olsman, Fangzhou Xiao, John C. Doyle

## Abstract

As we begin to design increasingly complex synthetic biomolecular systems, it is essential to develop rational design methodologies that yield predictable circuit performance. Here we apply theoretical tools from the theory of control and dynamical systems to yield practical insights into the architecture and function of a particular class of biological feedback circuit. Specifically, we show that it is possible to analytically characterize both the operating regime and performance tradeoffs of a sequestration feedback circuit architecture. Further, we demonstrate how these principles can be applied to inform the design process of a particular synthetic feedback circuit.

## 1 Introduction

A present challenge in synthetic biology is to design circuits that not only perform a desired function, but do so robustly. The difficulty in doing this arises in large part from the enormous amount of variability between both intracellular and extracellular environments [1–5]. While it is becoming easier to quickly implement a given circuit architecture [6,7], ensuring that its performance is robust to parameter variations is still a time-consuming and challenging task [8]. As synthetic circuits grow in size and complexity, it will be essential to develop a rational design methodology that allows biological engineers to easily identify the important design constraints for a given circuit and determine whether or not their desired behavior is feasible [9]. We can draw inspiration from the study of natural biological circuits, where cells are frequently confronted with a large amount of variability, yet exhibit robust behavior at the system level [10–16].

In the design of electrical and mechanical systems this problem is often solved with the implementation of feedback control [17,18], where the dynamics of a process are adjusted based on measurements of the system’s state with the aim of achieving some performance goals. For example, in a commercial cruise control system an engineer may want the car to be able to rapidly track whatever reference speed the user desires by measuring the car’s current velocity, while accelerating at a safe rate and not being too sensitive to small disturbances (e.g. road conditions). Similarly, a biological engineer may want the output concentration of a molecular species in a circuit to track an input signal (e.g. an inducer) with dynamics that are robust to parametric variability in reaction rates and the inherent noisiness of chemical kinetics [19].

Our primary focus here is a circuit architecture proposed by Briat et al. [20] that uses a sequestration mechanism to implement feedback control in a biomolecular circuit. This circuit immediately had a broad impact on the study of biological feedback systems, as sequestration is both abundant in natural biological contexts [21–23] and appears to be feasible to implement in synthetic networks [24–28]. For example, sequestration feedback can be implemented using sense-antisense mRNA pairs [29], sigmaantisigma factor pairs [30], or scaffold-antiscaffold pairs [19].

The purpose of this circuit is to control a process, composed of the molecular species *X*_1_ and *X*_2_ with two control species *Z*_1_ and *Z*_2_ (figure 1A). The goal is to set an external reference *μ* and have the concentration of the output species *X*_2_ robustly track it (figure 1B). The key property of this circuit is that it is able to implement robust perfect adaptation. This means that, at steady state, the concentration of *X*_2_ will be proportional to the reference *μ* (specifically, *X*_2_ = *μ*/*θ*_2_ on average). Importantly, the steady-state value of *X*_2_ will be *independent* of every other parameter of the network, implying its steady-state behavior is robust. We will focus first on studying a deterministic ordinary differential equation model of the circuit:

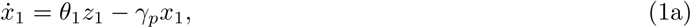

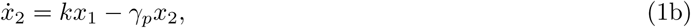

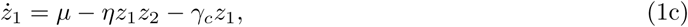

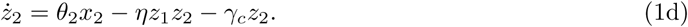

**Figure 1:**
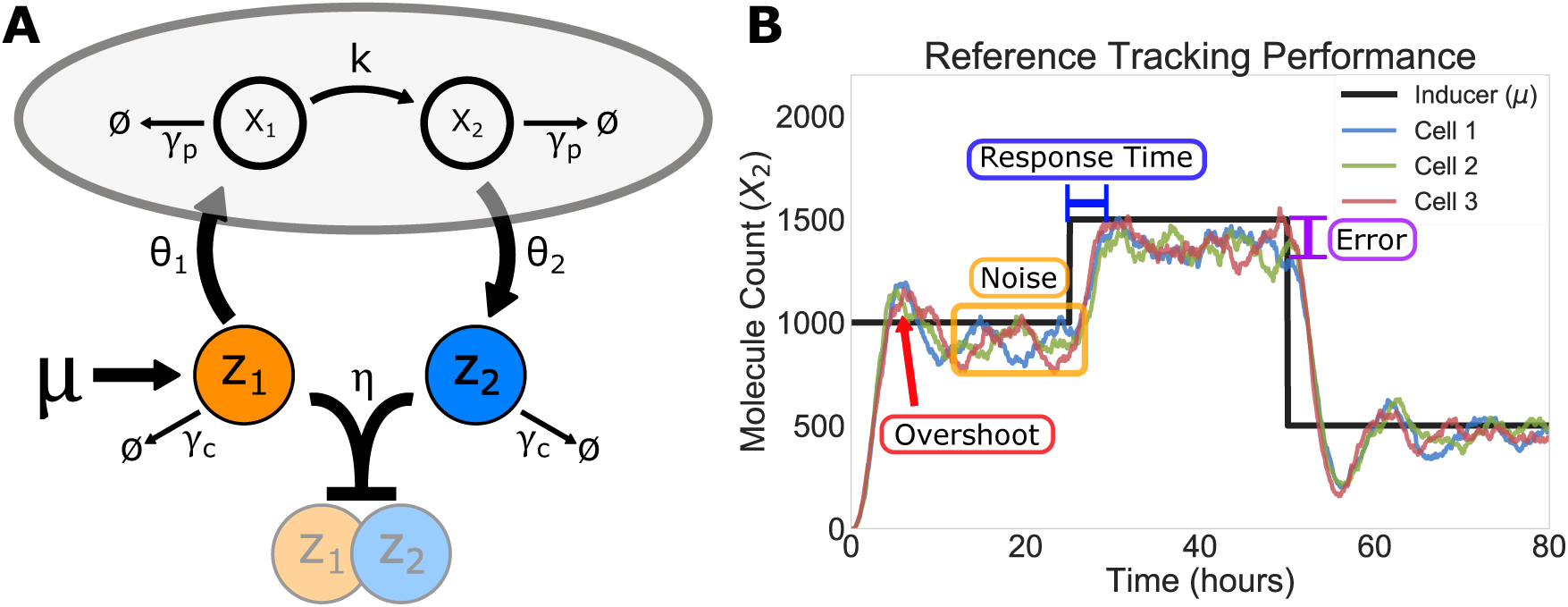
Characterizing Performance in the Sequestration Feedback Circuit. **A)** The circuit diagram for a class of sequestration feedback circuits, adapted from the model presented in [20]. Here we take *X*_1_ and *X*_2_ as the process species we are trying to control, and *Z*_1_ and *Z*_2_ as the controller species. One notable addition is that we explicitly model degradation of the control species *Z*_1_ and *Z*_2_ at the rate *γ_c_*. *θ*_1_ and *θ*_2_ represent the interconnection between the process species and the control species, *k* represents the *X*_1_-dependent synthesis rate of *X*_2_, and γ*_p_* is the degradation rate of the process species. Finally, *μ* acts as an external reference that determines production of *Z*_1_ through which we’d like to ultimately control the steady-state concentration of *Z*_2_, and *η* is the rate at which *Z*_1_ and *Z*_2_ irreversibly bind to each other. **B)** A representative plot of the type of behavior we expect from the circuit, where the concentration of *X*_2_ tracks a changing reference *μ*. We see that different cells have the same overall behavior, but with slight variations due to noise. This plot highlights the performance characteristics of this particular implementation of the circuit. For example we see that, when tracking the reference, *X*_2_ has some overshoot of the target (red), a period of time it takes to respond to changes (blue), random fluctuations due to noise (yellow), and steady-state error (purple). Ideally, we would like to a rational methodology to tune the circuit-level parameters of panel **A** to predictably control the system-level characteristics of panel **B**.

Here *θ*_1_, *θ*_2_, and *k* are production rates, *γ_p_* and *γ_c_* and degradation rates for the process species and controller species, respectively, *η* is the rate at which *η* and *z*_2_ sequester each other, and *μ* is the reference input that sets the synthesis rate of *z*_1_. We will refer to *x*_1_ and *x*_2_ as process species, and will focus in particular on *x*_2_ as the controlled output of the circuit. We can think of *z*_2_ as making measurements of *x*_2_, which are then propagated to *z*_1_ which can indirectly affect the production rate of *x*_2_ through *x*_1_. The use of lower-case letters for the species in equation (1) indicates that we are referring to a deterministic quantity (later in section 2.6 we will use upper-case variables to denote random variables).

If we take the viewpoint of an engineer designing this circuit for a particular application, for example controlling the expression of a downstream gene target, then we likely will have performance requirements that we would like the circuit to follow, such as responding to a change in reference within a given time frame and not transiently overshooting the reference by more than some set amount. The problem lies in understanding how these types of high-level system performance metric relate to the individual parameters of the circuit. Alternatively, if we consider natural circuits then we can take an evolutionary perspective, and study how different fitness constraints might shape the structure and function of existing biological feedback systems. Our work here uses methods from the field of control theory to shed light on this mapping between parameters and performance.

The structure and content of this article is somewhat unconventional, in that we present both a non-technical treatment of some of the core results in the companion piece [31], and also standalone research results that are not discussed in detail elsewhere. Where the content of [31] focuses on the application of classical tools in control theory to study the mathematical properties of the sequestration feedback circuit, the goal of this article is to outline practical guidelines that are accessible to an audience interested in utilizing these theoretical results to inform future experimental work in biological engineering.

In [20], the authors assume that the controller degradation rate *γ_c_* = 0 in equations (1c) and (1d), which yield the robust precise adaptation mentioned earlier. Our analysis of circuit performance and tradoffs in sections 2.1 to 2.3 makes the same assumption, and we address the case of *γ_c_* > 0 in section 2.4. In section 2.5 we apply this analysis to study a particular synthetic growth control circuit tha makes use of sequestration feedback. Finally, in section 2.6 we will investigate the effects of noise on the circuit.

## 2 Results

### 2.1 So Long As Sequestration Is Fast, It Can Be Ignored

The most obvious parameter to investigate in the sequestration feedback system is the sequestration rate *η*. To simplify our analysis, we will ignore controller species degradation for the time being (*γ_c_* = 0 in equations (1c) and (1d)), and analyze the effect of *γ_c_* in later sections. Because sequestration is a bimolecular interaction, *η* is a rate of association and has units of the form nM^−1^ h^−1^. Since the binding of *z*_1_ and *z*_2_ at the rate *η* in equations (1c) and (1d) encapsulates feedback in the system, we know that *η* cannot be too small. If sequestration is sufficiently slow, it will be as if there is no feedback in the circuit at all. More generally, a small *η* corresponds to a slow feedback action that tends to lead to a large overshoot of the desired steady-state, as can be seen in figure 2A.

The question then becomes, how large should *η* be? Briat et al. in their original work on this system observed that some sets of parameters lead to unstable behavior (which we discuss in greater depth in section 2.2), so it is important to investigate whether or not a large *η* could ever destabilize the circuit [20]. Ideally, we would like the find a regime of parameters where the system’s behavior is easy to predict and we do not have to worry about fine-tuning *η*. While it would be possible to get a sense for the behavior of *η* by simulating a broad parameter sweep and analyzing the resulting dynamics, we find that it is possible to gain a precise understanding of effects of *η* via theoretical analysis (described in detail in [31]).

Before we can analyze the effects of the sequestration rate on the circuit, we first need some notion of to what quantity it even makes sense to compare *η*. If nothing else, this quantity must have the same units as *η*, which immediately rules out a direct comparison with any other rate parameter in equation (1), as *η* is the only association rate in the system. Since *μ* is the only other parameter that has units of concentration (specifically nMh^−1^), it must be involved in the comparison. Dimensional analysis can get us as far as noting that a quantity of the form *α*^2^/*μ*, with *α* taking units of h^−1^, would at least have same units as *η*. This is as far as dimensional analysis alone can take us, since the other parameters in the system (*θ*_1_, *θ*_2_, *k*, and γ*_p_*) are all rates that have units consistent with *α*.

In [31], we find that *α* should take the form

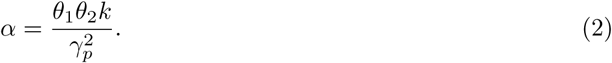

**Figure 2:**
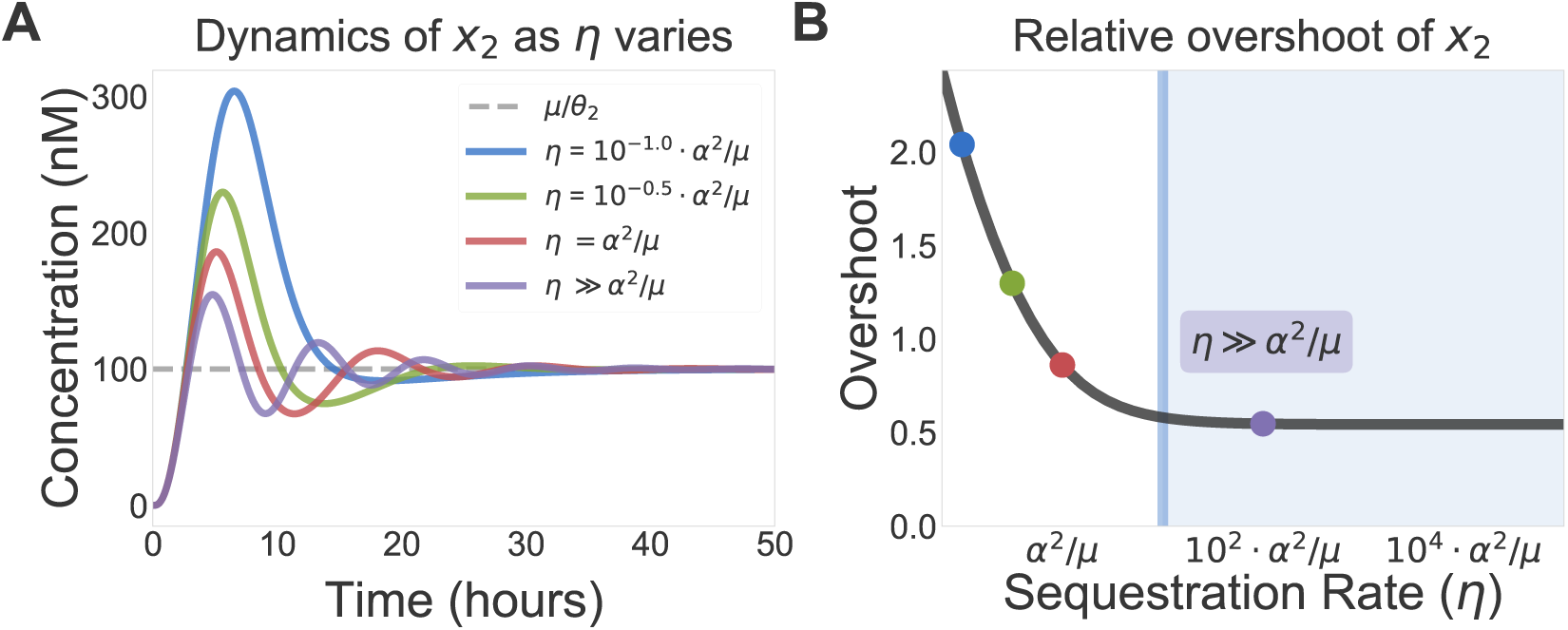
The Effects of Sequestration Rate on Dynamics. **A)** Here we show simulations of the circuit in equation (1) with *k* = *θ*_1_ = *θ*_2_ = γ*_p_* = 1h^−1^, *γ_c_* = 0 h^−1^, and μ = 100 nMh^−1^. This leads to a value of *α*^2^/*μ* = 10^−2^ nM^−1^ h^−1^. We vary *η* over the range 10^−3^ − 1 nM^−1^ h^−1^ and see that for small *η* the system’s response is highly sensitive on the sequestration rate. Once *η* is in the regime described by inequality (3), its dynamics are independent of *η*. **B)** A parametric plot showing this phenomenon, using the overshoot of the desired steady state 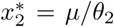 as a proxy for system performance. The black curve is generated by a simulated sweep of *η* values over the range 10^−3^ − 1 × 10^3^ nM^−1^ h^−1^, demonstrating that the overshoot becomes almost entirely invariant to *η* in the blue region where *η* > 10 · *α*^2^/*μ*. The colored dots correspond to the parameters of the simulations in **A**.

While the proof of this results is somewhat technical, we can derive from it a simple guideline for what it means to have *η* be large enough. The aggregate quantity α describes the steady-state rate at which *z*_2_ molecules are produced relative to the concentration of *z*_1_. In control theory, we would describe α as the *open-loop gain* of the system, since it captures the amplification between input and output in the absence of feedback. Characterizing the circuit in terms of α may yield practical benefits, as it allows us to sidestep the problem of individually measuring each of the four rate parameters in equation (2). We will see in later sections that many important features of the circuit can be written either in terms of the individual parameters in equation (1), or in terms of α.

Combining these ideas, we can now state precisely both what it means for *η* to be large and what effect this has on the circuit. We find that, though the behavior of the circuit is highly dependent on the value of all other parameters in the network, it is surprisingly insensitive to *η*, so long as

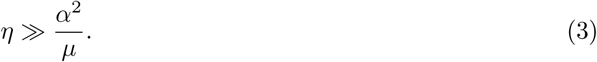

This inequality characterizes a separation of timescales between the production and degradation dynamics of the system (captured by *α* and *μ*), and the sequestration reaction. Intuitively, so long as sequestration is sufficiently fast, it does not affect the stability and performance of the circuit. This is demonstrated in figure 2A and B, where the system’s response becomes independent of *η* when inequality (3) holds. Here we use the amount of relative overshoot of *x*_2_ (defined as 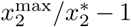) as a proxy for characterizing the circuit’s behavior. By simulating the circuit with *α*^2^/*μ* = 10^−2^ nM^−1^ h^−1^ as *η* varies logarithmically between 10^−3^ − 10^3^ nM^−1^ h^−1^, we see in figure 2B that the behavior of the circuit is insensitive to *η* when it is sufficiently large. The vertical line corresponds to *η* =10 · *α*^2^/*μ*.

The circuit’s stability and performance are also robust to changes in the *z*_1_ synthesis rate *μ*. Additionally, we observe that the circuit should be functional for any biologically plausible value of *μ*. One consequence of inequality (3) is that the steady-state values of *z*_1_ and *z*_2_ (denoted by *) must be have the relationship 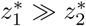. If measuring and comparing the rates in inequality (3) is not feasible, it is possible to tell if a circuit described by equation (1) is in the regime of strong feedback by simply comparing the concentrations of *z*_1_ and *z*_2_ at steady state.

We should emphasize that the circuit can still be functional when inequality (3) does not hold, however our analytic techniques yield less insight into what should be expected from the circuit in this regime. It is also important to note that inequality (3) is not sufficient to guarantee *good* performance (a concept into which we will delve more deeply in later sections). It merely implies that, once *η* is sufficiently large, the qualitative behavior of the circuit (good or bad) will not be affected by varying it. In the next sections, our analysis will focus on the parameter regime where inequality (3) holds and study how the rest of the rates in equation (1) affect the system’s dynamics.

### 2.2 Instability Arises from Production Outpacing Degradation

A central question for all feedback systems is whether or not the closed-loop circuit is stable. In many engineering applications, it can be the case that poorly implemented feedback control can destabilize an otherwise stable process [17]. This effect is quite salient in the sequestration feedback circuit, which was shown to have unstable oscillatory dynamics (known as limit cycles) for some parameter values. Before we can begin to consider how well a given set of parameters perform, we first need a guarantee that the corresponding dynamics have the baseline functionality of being stable. Ideally, we would be able to find system-level constraints that determine a priori when the circuit will be stable or unstable. While there exist straightforward numerical methods to predict whether or not a system with a given set of parameters is stable, it is substantially more difficult to derive general parametric conditions that characterize stability.

**Figure 3:**
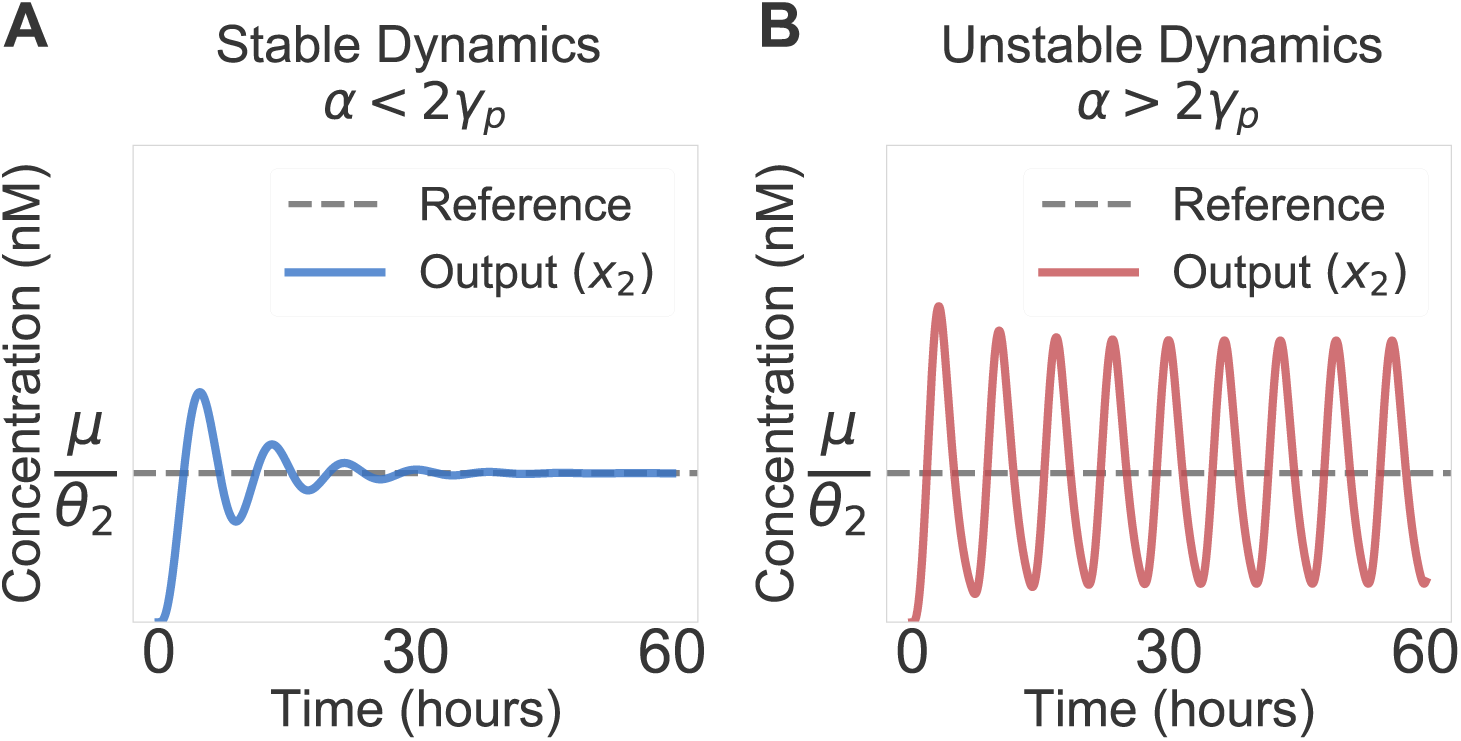
Dynamics can be Either Stable or Unstable. Here we demonstrate how inequality (4) affects the dynamics of the circuit in equation (1). **A)** This simulation uses *k* = *θ*_1_ = *θ*_2_ = γ*_p_* = 1h^−1^, *γ_c_* = 0h^−1^, *μ* = 100nMh^−1^, and *η* = 10nM^−1^ h^−1^. We see that *x*_2_ shows some transient oscillatory behavior, but ultimately adapts the the steady-state value *μ*/*θ*_2_. since a = 1h^−1^ and γ*_p_* = 1h^−1^, inequality (4) holds and the system is stable. **B)** Now we run the simulation with the same parameters, except *θ*_1_ = 3h^−1^. This implies that *α* = 3h^−1^, which tells us that inequality (4) no longer holds and the system will become unstable. In this case, the instability takes the form of indefinite oscillations. This figure is adapted from one presented in [31].

Take, for example, the relatively simple circuit described in equation (1), which has 7 free parameters. While it is straightforward to numerically simulate the circuit and investigate different parameter regimes, it is not at all obvious at first glance how the circuit will perform for a particular set of parameter values. We find that, in the limit of strong feedback and no degradation of *z*_1_ and *z*_2_, there exists a surprisingly simple relationship that determines stability. Using the same notation as in section 2.1, we find that the system is stable if and only if *α* < 2γ*_p_* (as shown in figure 3). This says that we need the net production rate in the circuit α to be slower than twice the degradation rate γ*_p_*. We can rewrite this result, using the fact that 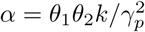, in the form

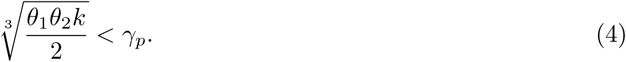

We note that the left-hand term is proportional to the geometric mean of all the production rates in the circuit. This is another perspective from which we can observe that production needs to be, on average, slower than degradation.

An interesting property of this inequality is that *μ* and *η* are conspicuously absent. The lack of dependence on *η* echoes the results we discussed in section 2.1, where we showed that the system’s performance is independent of the sequestration rate in the limit of large *η*. The fact that the system does not depend on *μ* makes sense because it is the only production rate in the system with units of concentration, so it must set the concentration scale for the circuit. We should expect that simply changing the units of concentration in the model should not affect stability. It then follows that varying *μ* is analogous to changing the units of concentration, and consequently should not affect stability and performance. We also find that the system is *intrinsically* stable when there is only one process species in figure 1A, which we prove in [31]. We find that for the case of *n* > 2 species we can derive results analogous to inequality (4), implying that there is a qualitative difference performance between *n* =1 species and *η* ≥ 2 species.

To further investigate inequality (4), we will assume that both the process parameters (γ*_p_* and *k*) and the desired set point (determined by *μ*/*θ*_2_) are fixed. Since we assumed that we are in the regime where *η* is large enough to not matter, the only remaining parameter to tune is *θ*_1_. We can interpret *θ*_1_ as quantifying the strength of interaction between the control species *z*_1_ and the process species *x*_1_. In this sense, inequality (4) tells us that there is a limit on how strong the connection between the controller and the process can be. It is then natural to ask how varying *θ*_1_ affects the circuits performance. To do this, we first need to define the relevant system-level performance metrics.

### 2.3 There is a Performance Tradeoff Between Speed and Robustness

In the previous section we noted that, for a given set of parameters, there is a maximum value that *θ*_1_ can take such that the controller remains stable. Picking the rate *θ*_1_ to be fast will speed up the response of the circuit, however if there is variability in *θ*_1_ the circuit may inadvertently exceed the limit set by inequality (4) and consequently become unstable. In this context, one way to characterize a circuit’s robustness is to determine not only if it is stable, but also that there are no small parameter changes that would result in it becoming unstable. A robust circuit is one that is far from instability, whereas a fragile circuit will be very near the stability boundary in inequality (4).

These observations yield a tradeoff: in order for the response time of the circuit to be as short as possible *θ*_1_ should be fast, however this necessarily makes the circuit more fragile. This notion of robustness is distinct from robustness of steady state, which is more commonly studied in biological contexts [13,14]. While steady-state robustness is fairly easy to define, as it is simply the error between the actual steady state of a system and a desired steady state, this notion of fragility requires more sophisticated mathematical tools and is generally difficult to solve for analytically. We discuss these theoretical methods in detail in [31], where we show that a good approximation of the fragility of the system has the form

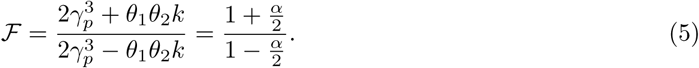

An equivalent interpretation of 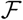 is as quantifying the system’s worst-case amplification of disturbances. As a sanity check we can see that, when we have equality in inequality (4) (i.e. 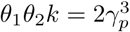), the fragility 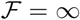 corresponding to the circuit becoming unstable. When 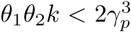, 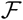 increases monotonically as *θ*_1_ increases. If *θ*_1_ = 0, then 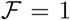 corresponding to no disturbance amplification (but also no control). We see in figure 4A that equation (5) yields a tradeoff curve that concisely capture the relationship between 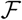 and θ_1_.

The upshot of this characterization is that we can now precisely quantify the performance tradeoff between speed and robustness. Where inequality (4) gave us a binary condition for stability, equation (5) provides a more nuanced measure of the circuit’s performance. Figure 4B demonstrates the effects of this tradeoff on the system’s dynamics. We see that, as the initial response time of the system decrease (*θ*_1_ increases), the system begins to oscillate and takes a much longer time to settle in to its steady-state value. These oscillations are indicative of the system approaching instability, a topic we explore in more depth in [31].

**Figure 4:**
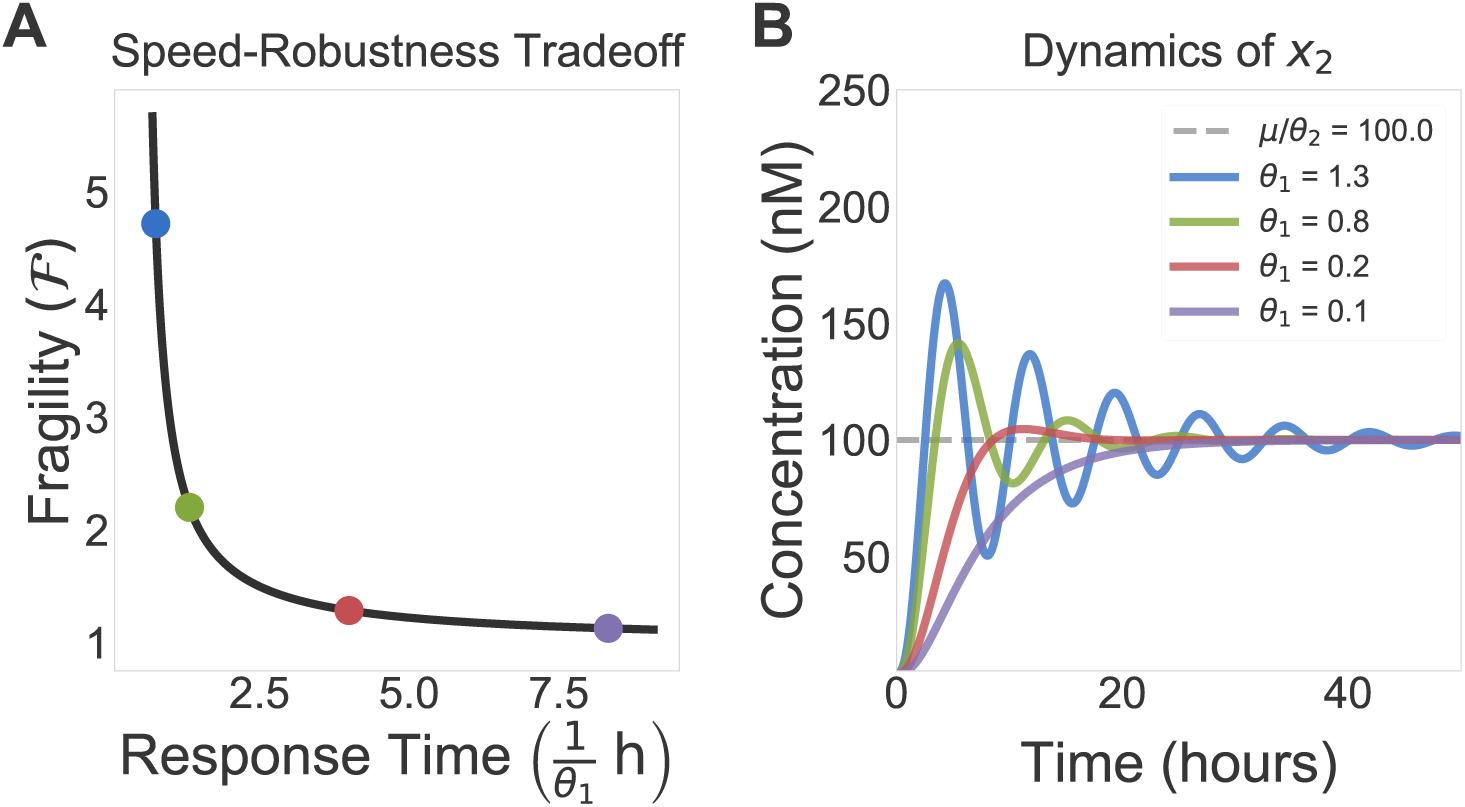
A Performance Tradeoff Between Response Time and Fragility. **A)** Here we show a tradeoff curve demonstration the relationship between response time (parametrized by 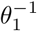) and fragility (as described in equation (5)). While an ideal system would have both a fast response and be minimally fragile (i.e. robust), this curve shows that, given all other parameters the system only has so much freedom to simultaneously optimize its performance. **B)** These trajectories corresponding to parameters associated with the colored dots in **A**, showing how fragility and response time relate to the actual dynamics of the circuit. We see that the blue curve rises quickly, but is highly oscillatory and takes a long time to settle. Conversely, the purple curve has a slow rise time, but is quite robust and settles quickly. In each plot we vary *θ*_1_ and use *k* = *θ*_2_ = γ*_p_* = 1 h^−1^, *γ_c_* = 0h^−1^, *μ* = 100 nMh^−1^, and *η* = 10nM^−1^ h^−1^. This figure is adapted from one presented in [31].

### 2.4 Controller Degradation Improves Stability but Introduces Steady-State Error

So far we have neglected the effects of degradation on the control species *z*_1_ and *z*_2_, assuming that they are only removed from the system by the sequestration reaction. Under this assumption, the results of the previous section tells us that the system is, in a sense, fundamentally constrained in its performance. We might hope, then, that nonzero controller degradation (*γ_c_* > 0) might give us an addition knob to tune in designing sequestration feedback circuits.

This turns out to be precisely the case, as is shown in figure 5. If all other parameters are held constant, the control species degradation rate *γ_c_* decreases the fragility of the system at the cost of introducing steady-state error. Again assuming the limit of large *η*, we show in [31] that there is a simple expression for this error, which we denote by e:

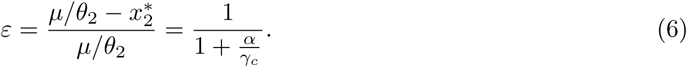

We can think of ε as capturing the steady-state error of *x*_2_ relative to the desired set point we would expect in the absence of controller degradation (*μ*/*θ*_2_). It is useful to think of controller degradation as adding ‘leakiness’ into the feedback mechanism. Imagine a scenario where *x*_2_ > *μ*/*θ*_2_. Ideally, this would cause an increase in *z*_2_ that precisely compensates for the mismatch. This increase in *z*_2_ will reduce the amount of *z*_1_ via sequestration. If, however, *z*_1_ is also degraded, there will be two mechanisms through which *z*_1_ is removed, namely degradation and sequestration. If we imagine a scenario in which *γ_c_* is extremely large, then every *z*_1_ molecule would likely be degraded before having a chance to be sequestered by a *z*_2_ molecule. If this were the case, then *x*_2_ cannot possible be using feedback to track the set point *μ*/*θ*_2_, because there is effectively no way for the increase in *z*_2_ to affect the concentration of *z*_1_. In equation (6), we see that ε = 0 if 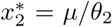 (no error), and ε = 1 when 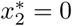 (maximum error).

**Figure 5:**
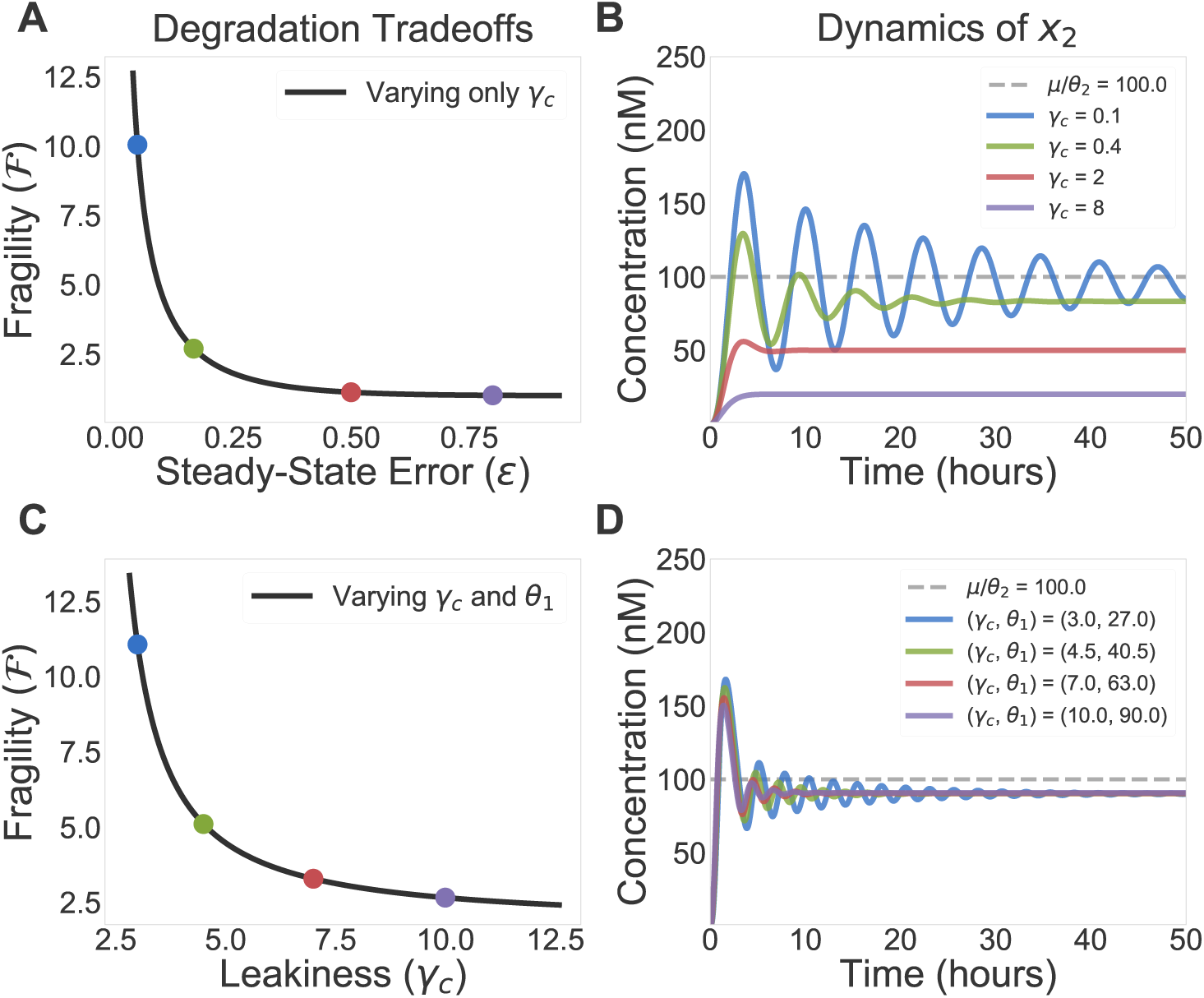
Controller Degradation Introduces Steady-State Error. **A)** We see that controller degradation *γ_c_* > 0 improves stability at the cost of introducing steady-state error. This tradeoff curve is a parametric plot where *γ_c_* is varied and equation (6) is compared to a generalization of equation (5) that incorporates the effects of *γ_c_*. **B)** Here we show the effects of the tradeoff in **A** on the dynamics of *x*_2_. The parameters are chose such that, if *γ_c_* = 0, the system would be unstable. The trajectory with small *γ_c_* (blue) is stabilized, but still has long-term oscillations indicative of fragility, but has little steady-state error. The trajectory with large *γ_c_* (purple) is extremely robust, but with large steady-state error. We vary *γ_c_* and and use *k* = 0_2_ = γ*_p_* = 1h^−1^, *θ*_1_ = 2h^−1^, *μ* = 100nMh^−1^, and *η* = 300nM^−1^ h^−1^. **C)** In this tradeoff curve, we hold the error ε = 0.1 constant by varying both *θ*_1_ and *γ_c_* as a constant ratio. We now observe that there is a tradeoff between fragility and leakiness, the latter parametrized by *γ_c_*. Intuitively, if *γ_c_* is large, then many copies of *z*_1_ and *z*_2_ are being degraded without ever being involved in the feedback process. **D)** We see that simulated trajectories display constant steady-state error, with less oscillatory behavior when *γ_c_* is large. Panels **C** and **D** use the same parameters as **A** and **B**, with the exception that *θ*_1_ is no longer fixed. This figure is adapted from one presented in [31].

It is also possible to describe the fragility 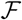 for the case where *γ_c_* > 0. This expression is much more complex than the one presented in equation (5), so we refer the reader to our analysis in [31] for details. The qualitative behavior of 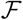 is shown in figure 5A, where increasing *γ_c_* decreases fragility. Combining these observations, we see that varying controller degradation introduces a tradeoff between steady-state error and robustness. Increasing *γ_c_* reduces 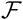 at the cost of increasing ε. Figure 5B demonstrates this effect via simulations of equation (1) with increasing values of *γ_c_*. We see that the trajectory with large controller degradation (purple) is far more stable than the trajectory with small degradation (blue), however the latter differs substantially from the original set point *μ*/*θ*_2_.

If it were the case that this error were simply a constant offset, then it could, in principle, be corrected for after the fact. The issue, however, is that the error is highly parameter dependent, as can bee seen from equation (6). If the goal is to ensure that the steady-state value of *x*_2_ is robust to variation in parameters, this error term may undermine the whole purpose of the circuit. Ideally, we would have some way to preserve the increased stability from controller degradation without suffering the consequences of large error. Fortunately, this is possible if we think not only in terms of varying *γ_p_*, but also the production rate *θ*_1_.

equation (2), tells us that *α* is proportional to *θ*_1_. This means that, if we increase *θ*_1_ to match an increase in *γ*_c_, it would be possible to hold the error in equation (6) constant. This would not necessarily be very interesting if this increase in *θ*_1_ increased 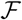 by a corresponding amount, however it turns out that we are still about to decrease 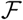 while keeping the ratio *θ*_1_/*γ*_c_ constant, as seen in figure 5C. We see in figure 5D this effect in simulations, where each trajectory has a constant steady-state error of ε = 0.1, however the trajectory with larger *γ_c_* values are significantly less oscillatory. This tells us that, if high turnover of *z*_1_ and *z*_2_ is not too costly, it is possible to mitigate the downside of degradation while preserving its benefits.

### 2.5 Sequestration Feedback in a Synthetic Bacterial Growth Control Circuit

The results presented so far have focused on the simple model of sequestration feedback presented in figure 1. This approach has facilitated our development of theoretical results that characterize some of the important features of sequestration as a mechanism for biological feedback control. We will now make use of the insights gained from the simplified model to study a particular synthetic bacterial growth control circuit. We show that the conceptual guidelines developed so far yield practical insight into the design of this circuit. Specifically, we find that the incorporation of controller degradation can lead to dramatically improved performance. The mathematical details of this analysis are presented in [31], here will show simulation results and explain at a high level how the theory can help guide circuit design.

A diagram of the circuit architecture is presented in figure 6A, where growth control is achieved by regulating the production of the toxin CcdB. Conceptually, if the intracellular concentration of toxin is proportional to the total number of cells, then the population as a whole will converge to a steady-state size that is less than the carrying capacity of the environment. The circuit uses a quorum sensing mechanism (involving the autoinducer AHL) to implement the coupling between population size and CcdB expression. The circuit described so far is capable of constant regulation, but lacks an extracellular mechanism through which we can control the population size (assuming that we do not want to be directly tuning protein expression, for example by altering the strength of ribosome binding sites).

The desired control can be implemented with molecular sequestration. For simplicity we will focus our modeling efforts on the particular mechanism of sense-antisense RNA pairing, however it would also be feasible to implement sequestration with sigma factor/anti-sigma factor binding or toxin/antitoxin binding. An antisense RNA (asRNA) is one that has a complimentary sequence to a messenger RNA (mRNA) strand. This complimentarity allows an asRNA to hybridize with an mRNA and block translation. By controlling the expression of asRNA, it is possible to modulate how responsive CcdB expression is to changes in AHL concentration and thus control the total size of the population. This architecture was originally proposed for experimental purposes in [32], and a functionally similar circuit was tested in [33]. We model this circuit with the following set of differential equations:

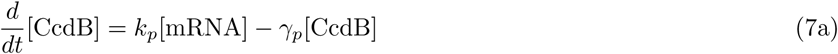

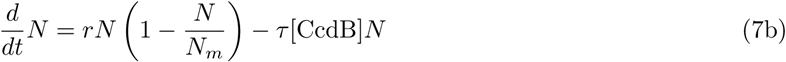

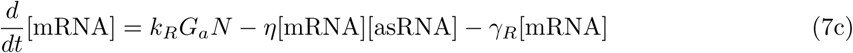

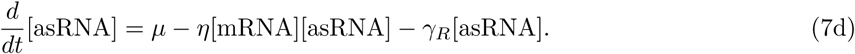

**Figure 6:**
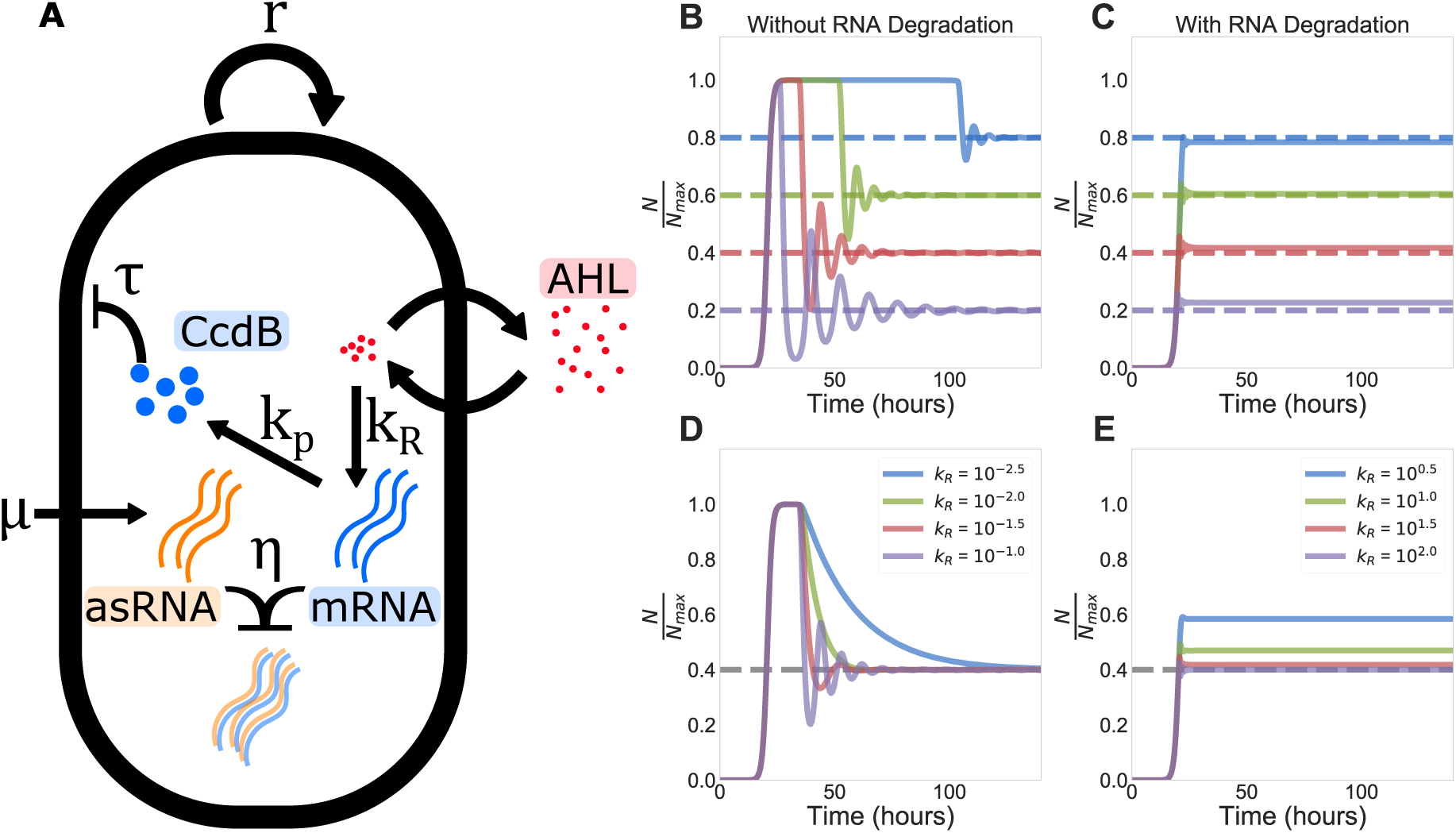
A Synthetic Growth Control Circuit. **A)** The circuit diagram for the dynamics described in equation (7). **B)** Simulations of the growth control circuit without RNA degradation (solid lines) for various set points *μ* (dashed lines). This architecture exhibits the precise adaptation property, though the response is relatively slow and oscillatory. **C)** Here we see the same circuit simulated with RNA degradation. The response is much faster and more robust, however there is non-zero steady-state error for each trajectory. **D)** Here we again simulate the circuit without degradation, but now vary *k_R_*. We see qualitatively similar performance tradeoffs to those in figure 4. **E)** As before, we see that adding controller degradation yields a very fast and consistent response. For these particular parameters, the circuit can achieve this performance with relatively little steady-state error. For all circuits we use the parameters *N_m_* = 10^9^, *r* = 1h^−1^, *η* = 20nM^−1^ h^−1^, *k_p_* = 10h^−1^, γ*_p_* = 3h^−1^, *G_a_* = 10^−6^ nM, and *τ* = 4 **×** 10^−3^ nM^−1^ h^−1^. Panel **B** uses *k_R_* = 0.1 h^−1^ and **C** uses *k_R_* = 10 h^−1^ and *γ_R_* = 20 h^−1^.

Quantities of the form [·] represent intracellular concentrations for each cell, and *N* represents the total number of cells. *N* follows logistic dynamics with an additional death rate due to toxicity τ proportional to the concentration of [CcdB] per cell. [CcdB] is a protein that is toxic to the cell, [mRNA] is the corresponding messenger RNA, the transcription of which we model as being induced by a quorum sensing ligand that is produced at a rate proportional to *N*, and [asRNA] is an antisense RNA that has a complementary sequence to the CcdB mRNA, thus acting as a sequestering partner. The term *G_a_* captures the gain between *N* and mRNA induction mediated by the quorum-sensing molecule AHL. We can think of [asRNA] and [mRNA] as representing *z*_1_ and *z*_2_, and the quantities [CcdB] and *η* as representing *x*_1_ and *x*_2_. This highlights the generality of the modeling framework in figure 1A: because we did not make any assumptions about the particular nature of the underlying variables, we are able to analyze a circuit with extremely heterogeneous underlying quantities (i.e. RNA, proteins, and cell population).

Figure 6B demonstrates how the growth control circuit adapts to various steady-state population levels when there is no controller degradation (*γ_R_* = 0). The steady state is set by varying *μ*. When possible, parameters for this model are taken from [34]. What is clear across all set points is that the population first grows to carrying capacity before the circuit is activated. Intuitively, the blue curve in Figure 6B has a large amount of asRNA that sequesters mRNA. Because of this, it takes longer to accumulate enough mRNA to make CcdB and lower the population level. In contrast, the purple curve has comparatively little asRNA, effectively increasing the rate at which CcdB can be produced. Qualitatively similar long-term oscillatory behavior in a CcdB-based growth control circuit was observed in [35].

Since [34] does not explicitly model transcription, we would ideally pick realistic transcription and translation timescales for bacteria. If we were to naively assume that we could model asRNA and mRNA as if they were like *z*_1_ and *z*_2_ in section 2.3, i.e. neglecting controller degradation and assuming they are only removed via sequestration, then we run into an issue. Because the sequestration mechanism modeled in section 2.3 assumes that controller degradation is negligible, we must use a very small mRNA synthesis rate to achieve stable dynamics, assuming all other parameters are fixed in a biologically plausible regime, even if sequestration is fast. This leads to a slow circuit response and a large transient overshoot. This is demonstrated in figure 6B, where CcdB production is so slow that the population reaches carrying capacity before the circuit can become active. In Figure 6D we see similar dynamics to those in figure 4, where the circuit faces harsh tradeoffs between speed and robustness.

We see from figure 6C and E that good performance requires not only that RNA is removed via sequestration, but also that it is degraded at a nontrivial rate. At the cost of a lack of precise adaptation, these circuits display dramatically improved performance. In figure 6C and E, the transient overshoot from figure 6B and D has almost entirely disappeared, and each system adapts on a nearly identical time scale, independent of *μ*. Figure 6D and ε compare performance for various values of *k_R_* in each system. Figure 6E shows that the introduction of *γ_R_* makes the system’s dynamics extremely robust to variations in *k_R_* over a wide range of values. We can interpret *γ_R_* as introducing a third tradeoff dimension, namely steady-state error. By allowing the system flexibility along this axis, its speed and robustness are greatly improved.

### 2.6 Noise and Fragility Are Two Sides of the Same Coin

Our analysis so far has assumed that the underlying circuit is perfectly deterministic, i.e. that its dynamics can be modeled by systems of ordinary differential equations. While these models serve as a good starting point for studying many biomolecular systems, they do not capture the effects of noise on the system. While noise is not always an important feature of biological processes, it can sometimes drastically alter the actual behavior of a circuit in a cell (for example, when certain molecules are at a low copy number) [5,36].

Here we will examine the steady-state variance of the output species *X*_2_ of the sequestration feedback system when its dynamics follow a stochastic chemical reaction model and relate it to the performance of the deterministic model equation (1). The capitalization in *X*_2_ reflects that this it is now a random variable, whereas *X*_2_ in equation (1b) is deterministic. To simplify our analysis, we will return to the assumption that there is no controller degradation (*γ_c_* = 0). In [20], the authors observed that there exist parameter values such that the deterministic model is unstable, but the average behavior of the stochastic model is stable (i.e. each species, on average, adapts to the desired steady-state concentration). We might think of each stochastic simulation representing a single cell’s dynamics, and the average representing the population-level behavior.

We find that, while this result regarding the mean behavior is correct, it does not tell the whole story. In particular we show that, while the population *average* is generally well behaved, the population *variance* can become large. In other words the population as a whole is predictable, however there is a large amount of cell-to-cell variability at any given time. In fact, it is the case that the noise scales in approximately the same way as the fragility of the system in equation (5) (as shown in figure 7A).

**Figure 7:**
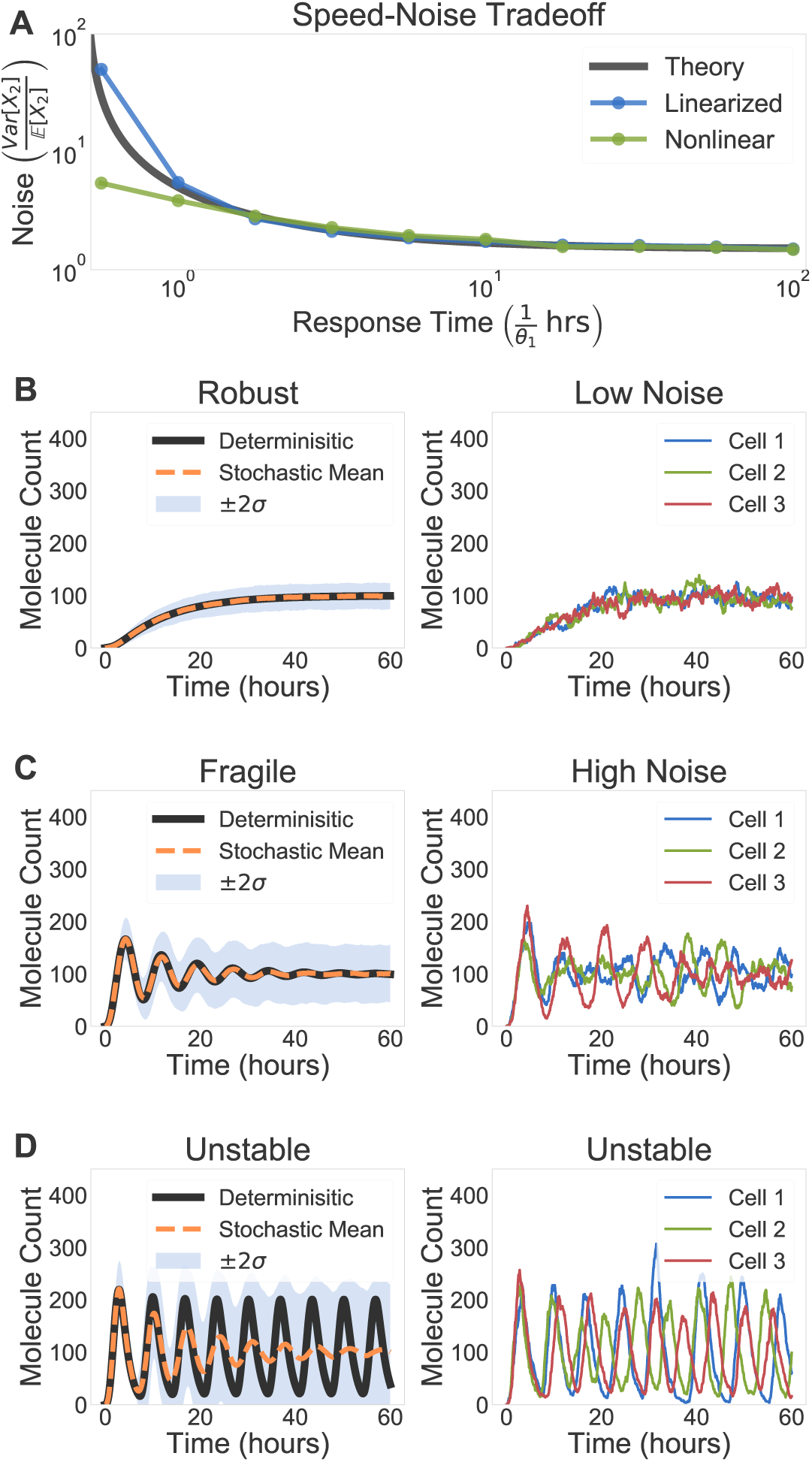
The Relationship Between Noise and Robustness. **A)** Here we see a general tradeoff between response time and noise (quantified by the Fano factor) of the sequestration feedback network. This is analogous to the tradeoff in figure 4A. The plot demonstrates the approximate behavior of equation (8) (black), simulation results for the same approximate model (blue), and simulation results for the fully nonlinear model without approximations (green). **B)** For *θ*_1_ = 0.75h^−1^, the deterministic and stochastic mean converge with good performance; individual stochastic trajectories are not very noisy. **C)** For *θ*_1_ = 1.3 h^−1^, the deterministic and stochastic mean have damped oscillations; individual stochastic trajectories are noisy. **D)** For *θ*_1_ = 3.5 h ^−1^, the deterministic model is unstable and oscillates while the stochastic mean is stable, as demonstrated in [20]. We see however, that the individual trajectories oscillate with randomized phase. In all simulations *k* = *θ*_2_ = *γ_p_* = 1 h^−1^, *η* = 10 h^−1^ nM^−1^, *μ* = 100 nMh^−1^. Mean trajectories and standard deviations are computed using *η* = 1000 trajectories.

Formally, we derive an approximate expression for the steady-state Fano factor (the variance divided by the mean) of *X*_2_ in the limit of fast sequestration:

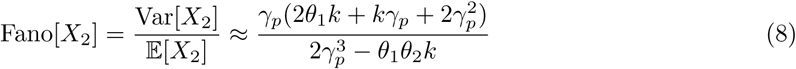

We see in figure 7A that the variability of *X*_2_ increases as the deterministic system approaches instability. This illustrates that there is a fundamental tradeoff such that the system can be either fast and noisy or slow and accurate, mirroring the deterministic tradeoff described in figure 4. We can get a sense for why this happens by observing that the denominator in equation (8) is the same as the denominator of equation (5). This tells us that we can expect each expression to grow in the same way as the respective denominators approaches 0. Thus, there is an intimate connection between the sensitivity of the deterministic sequestration feedback system, which corresponds to oscillatory behavior, and the sensitivity of the stochastic sequestration feedback system, which corresponds to increased noise. To give a more concrete sense for this relationship, we present representative simulation results that demonstrate this behavior.

In figure 7B, we see that a slow and robust deterministic performance (in the sense described in section 2.3) corresponds to a stochastic model with low noise. The left panel shows the mean behavior matching closely to the deterministic trajectory, with a fairly small amount of noise throughout the simulations. The right panel displays some sample individual trajectories, which essentially look like we would expect: closely following the mean with small deviations. If we look at figure 7C, we see that the deterministic model and the mean of the stochastic model converge to the reference quickly, with damped oscillations. Just as the fragility of the deterministic system is larger in the left panel, we see that the corresponding noise in the stochastic system is much larger than in figure 7B. Just as speed increased fragility in section 2.3, it appears to increase variability here. We note that these results assume that the sequestration reactions constitutes the only feedback in the system, recent work has shown that additional feedback loops can potentially serve to reduce noise [37].

Finally, figure 7D demonstrates a parameter regime where the deterministic model becomes unstable. In the left panel we see precisely the type of behavior described by Briat et al. in [20], where the stochastic mean appears to converge despite deterministic instability. The right panel, however, demonstrates that each individual trajectory is in fact exhibiting noisy oscillations, but with phases that are randomized relative to one another. Each individual cell is unstable, but this instability averages out at the population level. This highlights the importance of distinguishing between average and individual behavior.

## 3 Discussion

Though we could have, in principle, made some of the qualitative observation presented in this work from simulations alone, it is important to emphasize the fact that these theoretical results not only formalize numerical observations, but also force us to state exactly what it is we are measuring. An important contribution from [20] was not just that the authors proposed a clever mechanism to implement feedback control in biological contexts. They also went to great effort to clearly state and prove the existence of that the circuit is capable of achieving what they claimed. Our work here and in [31] is an attempt to pursue this line of reasoning and further characterize the qualitative and quantitative behavior of this circuit architecture.

More generally, this theoretical perspective sheds light on a variety of non-trivial parameter relationship which we hope will allow researchers avoid the need for brute-force parameter tuning when designing future control circuits. Where this article was intended to provide a relatively non-technical description of the work, the interested reader can find a great deal more mathematical depth and generality in [31]. We believe that this an exciting time for biology, where theory and experiment can productively guide each other towards new and interesting directions of inquiry.

## Author Contributions

Conceptualization and Methodology, NO, FX, and JCD; Formal Analysis, NO and FX; Software, NO, FX; Writing, NO; Supervision and Funding, JCD.

## Acknowledgements

The authors would like to thank Harry Nunns for providing feedback on the manuscript, Anandh Swaminathan for helping with stochastic simulations, and Reed McCardell for providing insight into the synthetic growth circuit. The project was sponsored by the Defense Advanced Research Projects Agency (Agreement HR0011–17–2–0008). The content of the information does not necessarily reflect the position or the policy of the Government, and no official endorsement should be inferred.

## 4 Supplemental Information

### 4.1 Analysis of Stochastic Systems

In this section we derive the approximate expression for the Fano factor of the output species in a stochastic sequestration feedback system with two process species, which was used in section 2.6. In the following we start from the chemical reaction network description of the sequestration feedback system [38–40], write down the chemical master equation for the stochastic system, and perform approximations with justifications to obtain an expression for the Fano factor of the output species. The approximation used is mathematically the same as the so-called linear noise approximation or first order system size expansion [36].

We describe the biochemical reactions of sequestration feedback system with two process species:

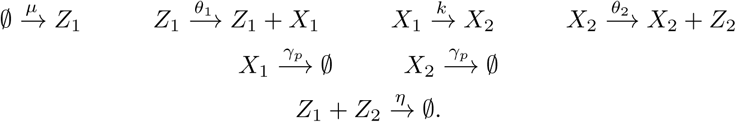

If we assume infinitely strong binding of the sequestration reaction, then limit *η →* ∞ holds. Hence, at any time, only one of species *Z*_1_ and *Z*_2_ can be non-zero. If both species are non-zero, then they sequester each other infinitely fast through reaction 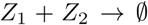 until one of them becomes zero. Therefore, we can define variable *Z* = *Z*_1_ − *Z*_2_, which has a one-to-one correspondence to species *Z*_1_ and *Z*_2_ counts, where positive Z indicates counts of *Z*_1_, and negative Z indicates counts of *Z*_2_.

With this simplification, the dynamics of the stochastic sequestration feedback system can be described by a continuous-time Markov chain (CTMC) over the counts of species *Z*, *X*_1_ and *X*_2_ using the following master equation dynamics [9]:

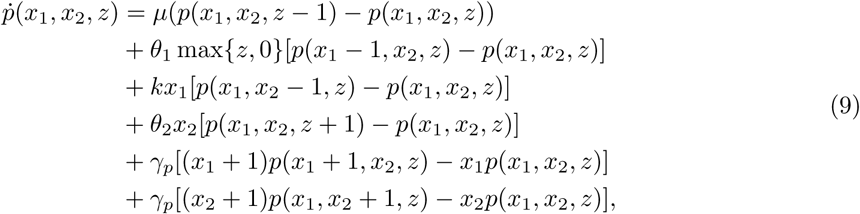

where *p*(*x*_1_, *x*_2_, *z*; *t*) denotes the probability for the system to have *Z* = *z*, *X*_1_ = *x*_1_, and *X*_2_ = *x*_2_ at time *t*.

We observe that all the terms on the right hand side of Equation (equation (9)) are linear, except for the max{*z*, 0} term. We can see this more clearly if we consider the first moment equation.

If we consider the steady state master equation, we set the left hand side to 0 and we apply 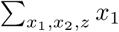 with the sum over all *x*_1_, *x*_2_ ∈ ℕ *z* ∈, then we obtain that

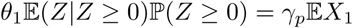

Similarly, if we apply 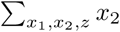 and 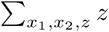

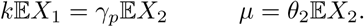

The term that prevents us from solving this set of linear equations for the first moments is the max{*z*, 0} term, which results in the probability for *Z* to be non-negative in the moment equations.

Therefore, we make a second assumption that *Z* ≥ 0 with probability 1 at steady state. This means *Z*_2_ is zero with probability 1 and this represents a good approximation if the system is stable, without *Z*_1_ oscillating to a very low count.

Under this assumption, we then obtain the linear equation:

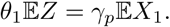

Similarly, if we apply sum 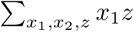 to the master equation, we obtain a system of linear equations for steady-state moments of both the first and the second order terms. As the system of equations becomes cumbersome to solve by hand, a Mathematica script was written to automatically derive and solve the moment equations. Solution gives the Fano factor of *x*_2_ as the following:

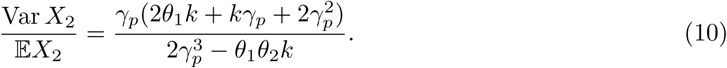

As *γ_p_* − ∞, we obtain an additional simplification

